# Deletion of competence genes represses expression of genes associated with anaerobic respiration/metabolism in *Aggregatibacter actinomycetemcomitans*

**DOI:** 10.1101/2023.05.18.541267

**Authors:** Nelli Vahvelainen, Laura Kovesjoki, Terhi Maula, Riikka Ihalin

## Abstract

Biofilm formation contributes to the virulence of various pathogens, as the extracellular polymer matrix provides protection against the host immune defense and antimicrobial drugs. Biofilm-associated diseases often become chronic and recurring. The periodontal pathogen *Aggregatibacter actinomycetemcomitans*, which resides in a multispecies biofilm in the subgingival pocket, produces multiple virulence factors that can contribute to disease progression. Certain strains of the species are naturally competent, which allows uptake of extracellular DNA that can be incorporated into the bacterial genome or used as a nutrient. Earlier studies indicated that bacterial interleukin receptor I (BilRI) and the type IV pilus subunit PilA protein are needed for efficient transformation in *A. actinomycetemcomitans*. In this study, we show that the outer membrane secretin HofQ is required for natural competence, as deletion of the *hofQ* gene results in a nontransformable strain. Furthermore, we studied the gene expression profiles of three single-gene mutants of the naturally competent *A. actinomycetemcomitans* strain D7S, all of which have decreased transformation efficiency compared to the wild-type strain. Additionally, as *A. actinomycetemcomitans* can bind to and internalize interleukin (IL)-1β, the effect of IL-1β on bacterial gene expression was also studied. However, in our experimental setup, the addition of IL-1β did not change gene expression in the *A. actinomycetemcomitans* strains used. The mutant strain lacking the *bilRI* gene exhibited a gene expression pattern similar to that of the wild-type strain. However, deletion of *hofQ* or *pilA* resulted in altered gene expression. Interestingly, genes associated with anaerobic growth, biofilm formation, and virulence were downregulated in the Δ*hofQ* and Δ*pilA* deletion mutants, which could indicate a decreased colonization ability and reduced virulence.

## Introduction

Microbial biofilms are associated with numerous chronic diseases in various anatomical locations, for example, endocarditis, cystic fibrosis, and periodontitis [1]. Moreover, biofilms can grow on medical devices such as catheters and mechanical heart valves [2]. Microbes embedded in biofilms are efficiently protected against the action of host immune cells and antimicrobial drugs by an extracellular polymer matrix. Thus, complete clearance of pathogenic microbes is difficult, and persistent cells can cause recurrent infections [3]. Natural biofilms typically consist of multiple species inhabiting different parts of the biofilm, as nutrients and gases are distributed unequally among the biofilm layers [4].

Periodontitis is a common oral inflammatory disease caused by a dysbiotic subgingival biofilm consisting mainly of gram-negative species. Progression of the disease can ultimately lead to tissue destruction and detachment of teeth [5]. The gram-negative opportunistic pathogen *Aggregatibacter actinomycetemcomitans* has been associated with periodontitis: along with other pathogenic species, it likely contributes to the shift from homeostasis to dysbiosis [6,7]. Although the gingival pocket is the conventional habitat of *A. actinomycetemcomitans*, this opportunistic pathogen can cause extraoral infections such as endocarditis, rheumatoid arthritis, and brain abscesses [8–10]. *A. actinomycetemcomitans* produces several virulence factors, such as RTX toxin family leukotoxins, cytolethal distending toxin (CDT), peptidoglycan-associated lipoprotein (PAL) and lipopolysaccharide (LPS) [11–14]. *A. actinomycetemcomitans* has also been shown to bind to and internalize human cytokines [15]. We previously showed that internalization of interleukin (IL)-1β alters the biofilm composition and decreases the metabolic activity of *A. actinomycetemcomitans* and that these changes are inhibited by deletion of either the outer membrane lipoprotein BilRI or the outer membrane secretin HofQ [16,17]. However, to our knowledge, no transcriptional studies have been performed with cytokine-treated *A. actinomycetemcomitans* biofilms.

Certain *A. actinomycetemcomitans* strains are naturally competent, which allows them to internalize extracellular DNA (eDNA) and incorporate it into their genome via homologous recombination [18,19]. The competent *A. actinomycetemcomitans* strains preferentially internalize DNA that contains a specific uptake signal sequence (USS) [20]. Several genes have been associated with natural competence, such as the type IV pilus gene cluster *pilABCD* and the regulator *tfoX (sxy)* [18,21,22]. Recently, we showed that the outer membrane lipoprotein BilRI also plays a role in efficient natural transformation [23]. The outer membrane secretin HofQ of *A. actinomycetemcomitans* is homologous to the competence protein ComE and exhibits structural and sequence properties similar to those of DNA-binding proteins [24]. Furthermore, the extramembranous part of HofQ (emHofQ) has been shown to bind DNA [24]. However, to date, no experimental studies have been performed to confirm the direct involvement of HofQ in the natural competence of *A. actinomycetemcomitans*.

In this study, we show that the outer membrane secretin HofQ of the naturally competent *A. actinomycetemcomitans* strain D7S is required for natural competence, as deletion of the *hofQ* gene resulted in a nontransformable strain. Furthermore, we studied how independent deletion of the competence-associated genes *bilRI*, *hofQ* and *pilA* affected the global gene expression profile of the naturally competent *A. actinomycetemcomitans* strain D7S. IL-1β was added to some cultures to study its effect on gene expression; however, the cytokine did not induce any changes in gene expression in the *A. actinomycetemcomitans* biofilms in our experimental setup. Comparison of gene expression profiles of the wild-type and deletion strains showed that the Δ*bilRI* mutant resembled the parental wild-type strain, but deletion of either *hofQ* or *pilA* resulted in changes in gene expression. The differentially regulated genes were associated with multiple functions, such as anaerobic respiration, anaerobic metabolism, carbohydrate metabolism, membrane biogenesis, transcription, and virulence. Overall, the changes in gene expression could indicate decreased biofilm formation and decreased virulence.

## Materials and methods

### Bacterial strains and growth conditions

The *A. actinomycetemcomitans* clinical strain D7S [25] and the deletion mutant D7S Δ*pilA::spe^r^* [21] were kind gifts from Prof. Casey Chen (University of Southern California, Los Angeles, CA, USA). The construction of the deletion mutants D7S Δ*bilRI* and D7S Δ*hofQ* has been described previously [16,17]. *A. actinomycetemcomitans* was grown in a candle jar at 37 °C.

### Natural transformation efficiency

The involvement of the outer membrane secretin HofQ in the natural transformation of *A. actinomycetemcomitans* strain D7S was studied by the natural transformation assay described in [20] with modifications described in [23] using the wild-type D7S strain and the deletion mutant D7S Δ*hofQ*. The construction of the linear DNA used in this assay (kanamycin resistance cassette flanked by sequences complementary to the genome of *A. actinomycetemcomitans* D7S) has been described in [23].

Briefly, plate-grown cells were collected in modified tryptic soy broth (mTSB: 30 g/l tryptone soy broth, 6 g/l yeast extract, 8 g/l glucose). Droplets containing 2×10^7^ cells were plated on tryptic soy agar (30 g/l tryptic soy broth, 3 g/l yeast extract, 15 g/l agar) supplemented with 5% heat-inactivated horse serum (hsTSA) and cultured for 2 h. The cells were then carefully mixed with 25 ng of the linear DNA construct using a metal wire loop, and the plates were incubated for 5 h, allowing the cells to take up the DNA. Sterile water without DNA was used as a negative control. After 5 h, the cells were collected in mTSB and plated on hsTSA supplemented with kanamycin (30 µg/ml). After 3–6 days, the transformant colonies were counted. The transformation efficiency was calculated as the number of colonies per microgram of added linear DNA. The transformation efficiency of the deletion mutant Δ*hofQ* was compared to the transformation efficiency of the wild-type strain. The assay was repeated eight times in triplicate.

### Bacterial cultures for RNA isolation

*A. actinomycetemcomitans* D7S, D7S Δ*bilRI*, D7S Δ*hofQ* and D7S Δ*pilA::spe^r^* were grown for 3 days on tryptic soy agar (37 g/l tryptone soy agar, 3 g/l agar) supplemented with 5% defibrinated sheep blood. The cells were collected in mTSB, and a suspension containing 0.5 × 10^9^ cells [26] in 5 ml of mTSB was transferred into a 50-ml cell culture flask (Cellstar® #690160, Greiner Bio-One, Germany). The cells were cultured for 20 h to allow them to attach to the bottom of the flask and form a biofilm. The biofilms were rinsed with RPMI 1640 medium (#R7509, Sigma) supplemented with 0.6 g/l L-glutamine (#G7513, Sigma) (hereafter referred to as RPMI) and incubated for 4 h in 5 ml of RPMI. To study the effect of IL-1β on bacterial gene expression, the medium was replaced with fresh RPMI supplemented with either 10 ng/ml IL-1β (ReliaTech, Wolfenbüttel, Germany) or an equivalent volume (5 µl) of sterile water, and the biofilms were grown for an additional 2 h. The cells were harvested from the flasks by scraping and collected in 700 µl of RNA*later*® Solution (#AM7020, Ambion). The cells were pelleted (6,000 × g, 10 min, 4 °C), resuspended in 200 µl of RNA*later*®, and stored at 4 °C for RNA isolation (max. 1 week). The culture was repeated three times in duplicate.

### RNA isolation and sequencing

Before RNA isolation, RNA*later*® was removed from the cell pellets by centrifugation (6,000 × g, 5 min, 4 °C), and the cells were washed with nuclease-free water (#AM9938, Ambion) and collected by centrifugation as described above. RNA was isolated from the cell pellets with a RiboPure™ Bacteria Kit (#AM1925, Ambion) following the manufacturer’s instructions with some modifications. Briefly, the cells were mixed with the supplemented phenol reagent and lysed by vortexing with zirconia beads (5 × 2 min, 2 min incubation on ice in between). The beads were removed by centrifugation (16,000 × g), and 0.3 volumes of chloroform (without isoamyl alcohol, #C2432, Sigma) were added to the recovered lysate. After 10 min of incubation, the resulting aqueous phase (top) was collected and further purified on a spin column. RNA samples were treated with DNase I for 2 hours at 37 °C to ensure that all remaining genomic DNA was digested. The removal of genomic DNA was verified by 16S rRNA PCR (primers in Table 1).

**Table 1.**
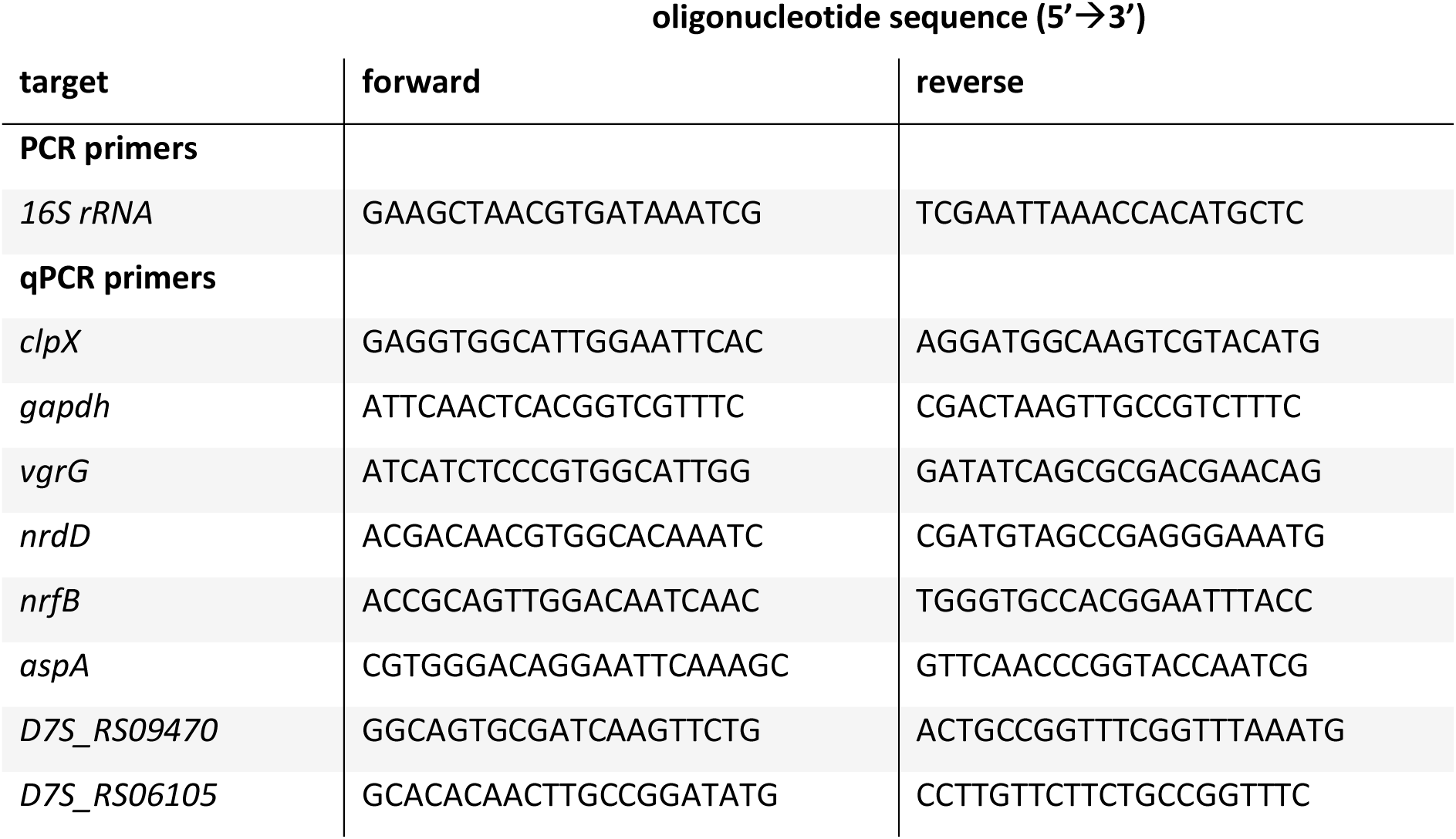
Primers used in this study.

The quality of the RNA samples was ensured using an Agilent Bioanalyzer 2100. The sample concentration was measured with Qubit®/Quant-IT® Fluorometric Quantitation (Life Technologies) and/or Nanodrop ND-2000 spectrophotometer (Thermo Scientific). Library preparation was performed according to the library preparation protocol (TruSeq® Stranded total RNA Reference Guide, Illumina (1000000040498)) with an Illumina Stranded Total RNA Prep Ligation with Ribo-Zero Plus Kit (Illumina). Sample quality was ensured using an Agilent Bioanalyzer 2100 or Advanced Analytical Fragment Analyzer, and the concentration was measured with the Qubit®/Quant-IT® Fluorometric Quantitation and/or KAPA Library Quantification kit for Illumina platform (KAPA Biosystems). Paired-end sequencing (read length 100 bp) was performed using Illumina NovaSeq 6000 S4 v1.5 instrument. Base calling was performed with bcl2fastq2 conversion software. RNA sample quality checks, library preparation and quality checks, and total RNA sequencing were performed at the Finnish Functional Genomics Centre (Turku, Finland).

### Data analysis

RNAseq data analysis was performed with CSC Chipster v.4 analysis software [27]. The quality of the raw data in all the fastq files was checked simultaneously with the MultiQC tool (based on the FastQC package [28] and MultiQC package [29]). The strandedness of the reads was checked with RseQC [30]. Due to the large number of reads per sample (90-160 M), subsets of 10 M reads were generated with a Chipster tool based on the seqtk package [31].

The subsets of reads were aligned to the *A. actinomycetemcomitans* strain D7S-1 genome (RefSeq accession NC_017846.2) with Bowtie2 [32]. The aligned reads were mapped to genes with HTSeq [33] using the genome annotation for *A. actinomycetemcomitans* strain D7S-1 (genome assembly ASM16361v3, RefSeq accession GCF_000163615.3). Differential expression analysis and principal component analysis were conducted with DESeq2 [34] with the cutoff for the Benjamini‒Hochberg adjusted p value set at 0.05.

Genes that were up- or downregulated at least 2-fold (log_2_-fold change ≥ 1 or ≤ 1, respectively) and had an adjusted p value of < 0.05 were selected for further analysis. Protein identifications (RefSeq Protein IDs) were retrieved from NCBI. The RefSeq Protein IDs were imported into the UniProt (release 2022-05) Retrieve/ID Mapping tool to generate Gene Ontology (GO) terms related to the proteins. Additionally, the RefSeq Protein IDs were searched for in the Clusters of Orthologous Genes (COG) database [35].

### Quantitative real-time PCR (qPCR)

The same total RNA samples were used for both RNAseq and qPCR. The relative gene expression levels of six selected genes (*nrfB*, *aspA*, *vgrG*, *nrdD*, D7S_RS09470, and D7S_RS06105) were measured with two-step RT‒qPCR using a Bio-Rad iCycler iQ5 real-time PCR system. Two reference genes (*clpx* and *gapdh*) were used to normalize gene expression levels. The sequences of the qPCR primers are presented in Table 1. Primer specificity was confirmed with melting curve analysis and/or agarose gel electrophoresis.

For each sample, 500 ng of total RNA was used as a template for cDNA synthesis with the LunaScript® RT SuperMix Kit (#E3010, New England Biolabs) following the manufacturer’s instructions. Synthesized cDNA and control RNA without reverse transcriptase (no-RT control) were diluted 1:5 in nuclease-free water. Dye-based qPCR was performed with Luna Universal qPCR Master Mix (#M3003, New England Biolabs) as recommended by the manufacturer: 1 µl of the diluted cDNA or no-RT control was added into a reaction mixture containing 10 µl of 2x Luna Universal qPCR Master Mix, 1 µl of each primer (5 µM) and 7 µl of nuclease-free water. The thermocycling protocol consisted of two cycles: Cycle 1 (1×)—Step 1: 95 °C for 60 s, Cycle 2 (40×)—Step 1: 95 °C for 15 s, Step 2: 60 °C for 30 s + plate read. Melting curves were generated by increasing the temperature from 60 °C to 95 °C in 0.5 °C increments with a 30 s incubation at each temperature. qPCR was performed with technical triplicates, including a nontemplate control for each primer pair. An interplate control sample containing 5 ng of the cDNA mixture as a template for amplification with the *gapdh* primers was included in triplicate in each plate to ensure reliable comparison of results between plates. Relative gene expression was calculated with a method developed by Pfaffl [36] using two housekeeping genes to normalize the expression levels [37,38]. Primer efficiencies were calculated from a 5-point cDNA standard curve generated with serial 2-fold dilutions. Wild-type samples were used as a control for relative gene expression.

### Correlation between RNAseq and qPCR data

The correlations between the relative gene expression values obtained by RNAseq and qPCR were analyzed with a linear regression model and Pearson correlation analysis (n = 12). The raw counts per gene obtained by RNAseq were normalized with DESeq2 (Chipster tool). Relative gene expression was calculated with respect to the wild-type samples. Similarly, gene expression values obtained by qPCR were compared to the average value of the wild-type samples. Pearson correlation coefficients were calculated with R (version 4.1.2) [39] using R Studio (version 2022.12.0.353) [40]. The output of the linear regression model was visualized with the ggpubr package [41].

### Statistics

Statistical analyses were performed with IBM SPSS (version 27) unless otherwise stated. Differences between two groups were analyzed with the nonparametric Mann‒Whitney U test due to the small sample size (n = 3–8). Differences were considered statistically significant at p < 0.05.

## Results

### The outer membrane secretin HofQ is required for natural transformation

Certain *A. actinomycetemcomitans* strains are naturally competent, which allows them to internalize extracellular DNA (eDNA) and incorporate it into their genome via homologous recombination [18,19]. Previous sequence and structural analyses have suggested that the DNA-binding outer membrane secretin HofQ of *A. actinomycetemcomitans* could be involved in transformation [24]. Therefore, we studied how deletion of the *hofQ* gene affected the transformation of the naturally competent *A. actinomycetemcomitans* strain D7S. Compared to the parental wild-type strain, the D7S Δ*hofQ* mutant had a transformation efficiency of 0–1.9% (p = 0.0002, Mann‒Whitney U test (n = 8)) (**Figure 1**). Moreover, 6 of the 8 experiments performed with the Δ*hofQ* mutant resulted in zero transformed colonies, strongly suggesting that the Δ*hofQ* mutant strain is nontransformable. Therefore, the results confirm that HofQ is required for natural transformation in *A. actinomycetemcomitans*.

**Figure 1.**
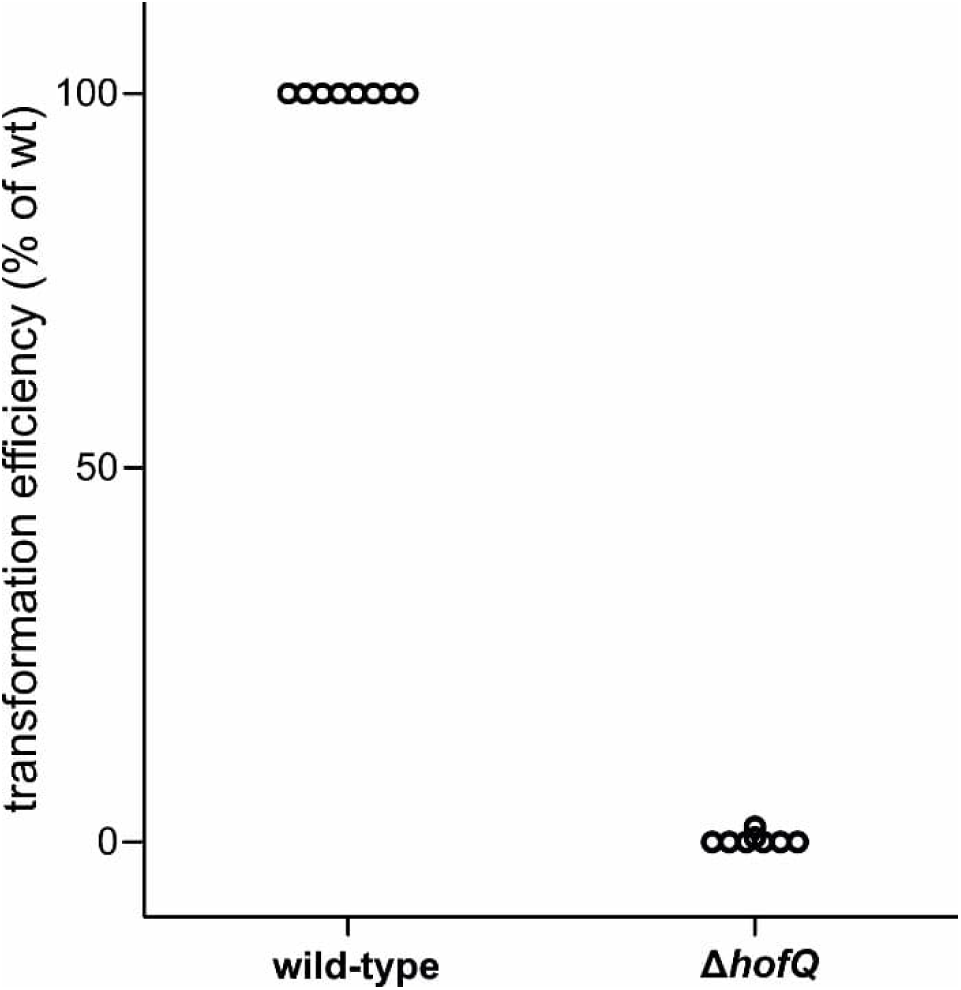
Deletion of *hofQ* in the naturally competent *A. actinomycetemcomitans* strain D7S resulted in a nontransformable strain (p = 0.0002, Mann‒Whitney U test (n = 8)). The transformation efficiency of the deletion mutant decreased to 0–1.9% of that of the parental wild-type strain, with a transformation efficiency of 0% in the majority of the experiments.

### IL-1β did not affect the gene expression profile of *A. actinomycetemcomitans* D7S

*A. actinomycetemcomitans* D7S has been shown to bind to and internalize IL-1β, which alters the biofilm composition and decreases the metabolic activity of the cells [15–17,42]. Deletion of the outer membrane lipoprotein BilRI decreases the internalization of IL-1β [16], and deletion of either *bilRI* or *hofQ* inhibits the IL-1β-induced changes in *A. actinomycetemcomitans* biofilms [16,17]. Therefore, we sought to investigate whether exposure to this cytokine affects bacterial gene expression.

Additionally, as the type IV pilus subunit PilA has been characterized as a potential virulence factor [43] and the deletion mutant Δ*pilA::spe^r^* has been shown to form rough phenotype cells similar to those of the wild-type strain [21], we sought to study the effect of IL-1β on the Δ*pilA::spe^r^* biofilm.

Biofilms of the wild-type *A. actinomycetemcomitans* D7S strain and the Δ*bilRI*, Δ*hofQ* and Δ*pilA::spe^r^* deletion mutants were incubated with 10 ng/ml IL-1β for 2 hours. Total RNA sequencing was performed for the IL-1β-treated and control cultures, but the results showed that the cytokine treatment did not alter gene expression in any of the strains. Principal component analysis showed some clustering among the different strains but not between the treatments (IL-1β/MQ) (**Figure 2A**). Because IL-1β did not affect the gene expression of *A. actinomycetemcomitans* in our experimental setup, we focused on the gene expression differences between the wild-type strain and each of the three deletion mutants. The IL-1β-treated samples were not included in the analysis.

**Figure 2.**
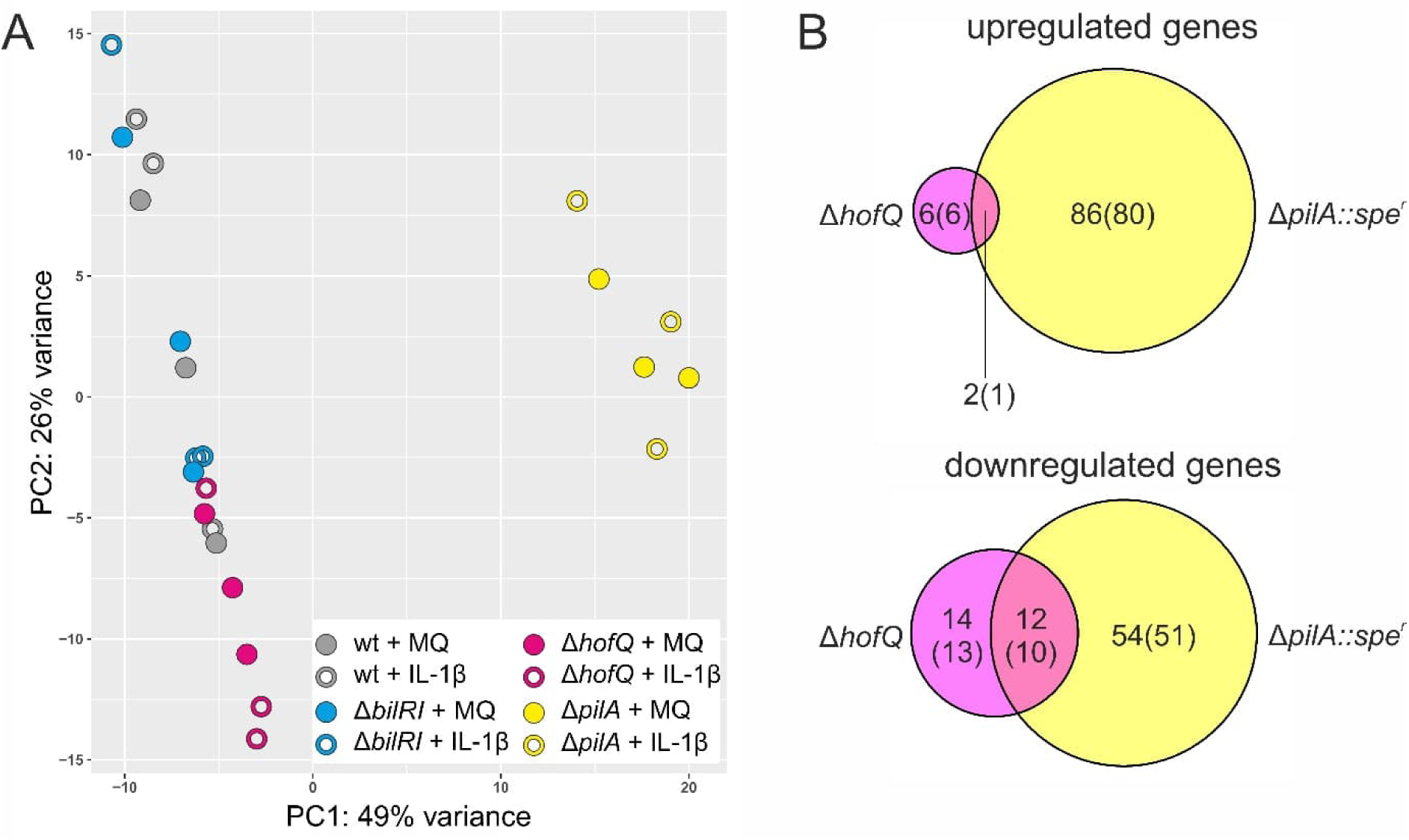
**A.** Principal component analysis of the total RNAseq data shows that while there was clustering among the different strains, there was no difference between the IL-1β-treated and control (MQ) cultures. **B.** In the Δ*hofQ* and Δ*pilA::spe^r^* strains, there were several genes whose expression level changed by at least 2-fold compared to that in the parental wild-type strain. The majority of the differentially expressed genes encode proteins (number in parenthesis).

### Deletion of *hofQ* or *pilA* changed the gene expression profile of *A. actinomycetemcomitans* D7S

Deletion of *bilRI* or *hofQ* changes the composition of *A. actinomycetemcomitans* D7S biofilms [16,17]. Additionally, deletion of *hofQ* significantly decreases biofilm mass [17]. Therefore, we sought to study whether deletion of these genes results in significant changes in the transcription of genes. Furthermore, as both *bilRI* and *hofQ* are associated with natural competence [23], we sought to study how the deletion of an additional competence gene, *pilA* [21], affects gene expression in the naturally competent *A. actinomycetemcomitans* D7S strain.

Differential expression analysis with DESeq2 was used to identify differentially expressed genes between the wild-type strain and each of the three deletion mutants (Δ*bilRI*, Δ*hofQ* and Δ*pilA::spe^r^*). A threshold of a 2-fold change in gene expression was used to identify differentially regulated genes. As expected, the gene with the greatest negative fold change in each deletion mutant was the corresponding deleted gene. Otherwise, the expression profile of the Δ*bilRI* deletion mutant strongly resembled that of the parental wild-type strain, as the only gene whose expression was significantly different was *bilRI*. However, deletion of *hofQ* or *pilA* resulted in changes in overall gene expression. Compared to the parental wild-type strain, the Δ*hofQ* mutant had 26 downregulated and 8 upregulated genes, whereas the Δ*pilA::spe^r^* mutant had 66 downregulated and 88 upregulated genes (**Figure 2B**). The Δ*hofQ* and Δ*pilA::spe^r^* mutants shared 12 downregulated and 2 upregulated genes. The full list of differentially regulated genes in the Δ*hofQ* and Δ*pilA::spe^r^* mutants is presented in the supplemental information.

The majority of the differentially expressed genes in the Δ*hofQ* and Δ*pilA::spe^r^* mutant strains were protein-coding genes (**Figure 2B**). Thirty protein-coding genes were differentially regulated in the Δ*hofQ* mutant, and 142 were differentially regulated in the Δ*pilA::spe^r^* mutant. The two mutant strains shared 1 upregulated and 10 downregulated protein-coding genes. A total of 14 of the differentially regulated proteins between the two strains were hypothetical proteins. The differentially regulated proteins were categorized according to Gene Ontology (GO) terms. At least one GO term was found for most of the proteins: no GO terms were found for 6 proteins differentially regulated in Δ*hofQ* and 44 proteins differentially regulated in Δ*pilA::spe^r^*. GO terms are classified into three categories: biological process, cellular component, and molecular function. Differentially regulated proteins in both mutant strains were found in all three categories. GO enrichment analysis revealed that the molecular function term metal ion binding (GO:0046872) was one of the terms most enriched with both up- and downregulated proteins in Δ*hofQ* and Δ*pilA* (**Figure 3**A-B). Metal ions are common cofactors for a variety of enzymes, such as oxidoreductases, transferases, and ligases [44]. Another enriched molecular function term was ATP binding (GO:0005524). The most enriched cellular component terms cytoplasm (GO:0005737) and plasma membrane (GO:0005886) indicate the subcellular location of the regulated proteins.

**Figure 3.**
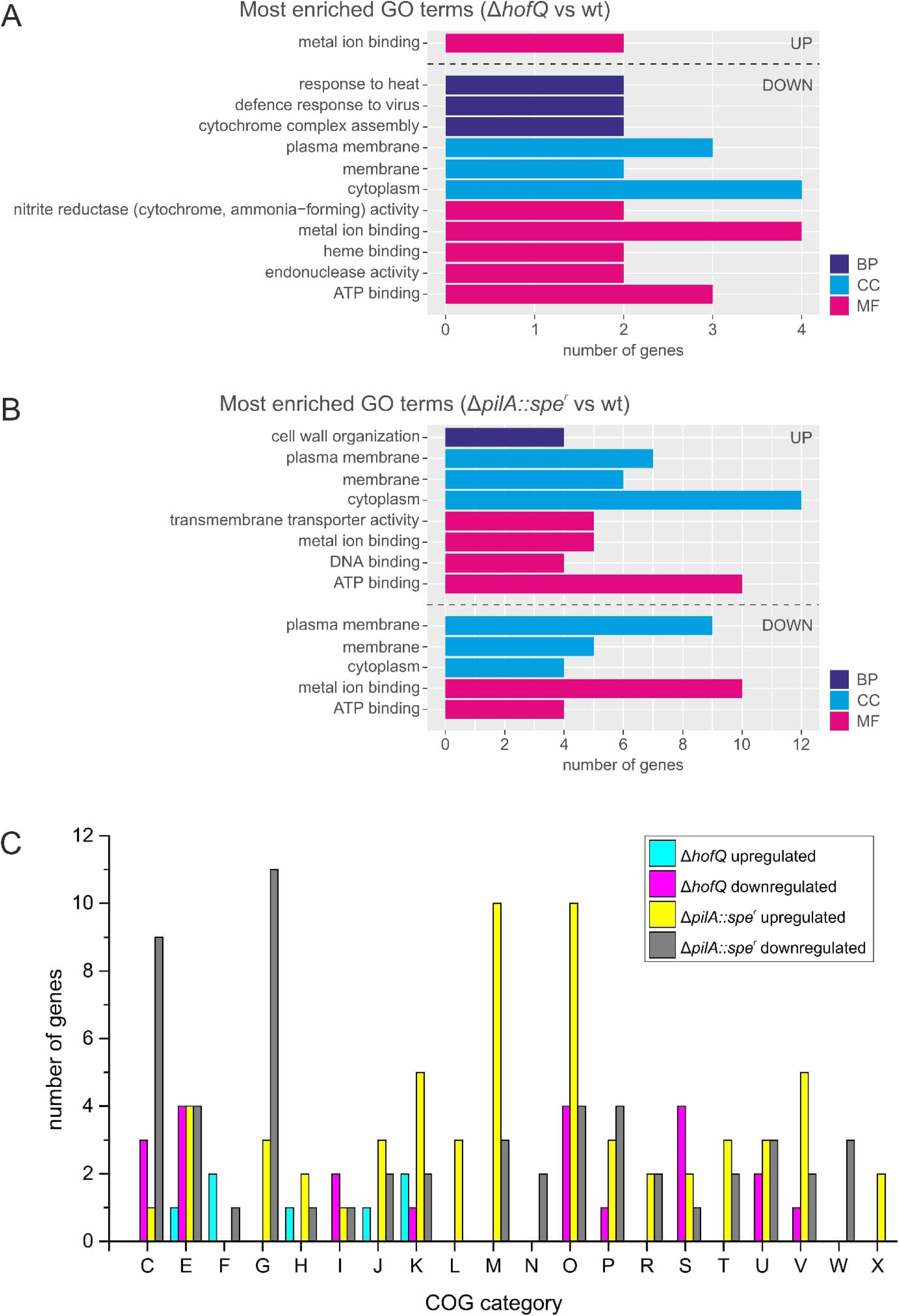
Classification of the differentially regulated proteins in the deletion mutants Δ*hofQ* and Δ*pilA::spe^r^*. Gene Ontology (GO) terms are classified into three categories: biological process (BP), cellular component (CC) and molecular function (MF). **A.** GO terms associated with ≥2 up- or downregulated proteins in the Δ*hofQ* deletion mutant. **B.** GO terms associated with ≥4 up- or downregulated proteins in the Δ*pilA::spe^r^*deletion mutant. **C.** Distribution of differentially regulated genes in Cluster of Orthologous Genes (COG) categories. C: energy production and conversion, E: amino acid metabolism and transport, F: nucleotide metabolism and transport, G: carbohydrate metabolism and transport, H: coenzyme metabolism, I: lipid metabolism, J: translation, K: transcription, L: replication and repair, M: cell wall/membrane/envelope biogenesis, N: cell motility, O: posttranslational modification, protein turnover, chaperone functions, P: inorganic ion transport and metabolism, R: general functional prediction only, S: function unknown, T: signal transduction, U: intracellular trafficking and secretion, V: defense mechanisms, W: extracellular structures, X: mobilome: prophages, transposons.

Additionally, the differentially regulated proteins were assigned to COG categories (**Figure 3C**). A category was found for most of the proteins: no category was found in the COG database for 4 differentially regulated proteins in Δ*hofQ* and 39 differentially regulated proteins in Δ*pilA::spe^r^*. Among the downregulated proteins in Δ*hofQ,* the most common COG categories were E (amino acid metabolism and transport), O (posttranslational modification, protein turnover, chaperone functions) and S (function unknown), while the upregulated proteins were nearly equally distributed among categories E, F (nucleotide metabolism and transport), H (coenzyme metabolism), J (translation) and K (transcription). In Δ*pilA::spe^r^*, proteins belonging to categories M (cell wall/membrane/envelope biogenesis) and O were upregulated, whereas proteins belonging to categories C (energy production and conversion) and G (carbohydrate metabolism and transport) were downregulated.

### qPCR validation of the RNAseq results

The expression of five genes that were differentially expressed in both Δ*hofQ* and Δ*pilA::spe^r^* was further analyzed by qPCR to validate the RNAseq results. Four of these genes (D7S_RS05920 *aspA*, D7S_RS09095 *nrfB*, D7S_RS06105 and D7S_RS09470) were downregulated and one (D7S_RS12360 *vgrG*) was upregulated compared to the parental wild-type strain. Additionally, one more gene (D7S_RS03435 *nrdD*) that was upregulated in the Δ*hofQ* mutant was selected for qPCR. None of the six selected genes were differentially expressed in the Δ*bilRI* mutant, according to the RNAseq results. Two housekeeping genes, *clpX* and *gapdh*, were selected for normalization of gene expression levels. The *clpx* gene (ATP-dependent Clp protease ATP-binding subunit ClpX, D7S_RS07855) was selected because it is constitutively expressed in *A. actinomycetemcomitans* biofilms [45]. The *gapdh* gene (glyceraldehyde-3-phosphate dehydrogenase, D7S_RS08265) has traditionally been used as a reference gene [46,47].

The qPCR results confirmed that among the three mutant strains, Δ*bilRI* had the most similar expression profile to the wild-type strain. Although the RNAseq results indicated that only *bilRI* was differentially regulated, the qPCR results showed that *aspA* was significantly downregulated in the Δ*bilRI* mutant (p = 0.0495, Mann‒Whitney U test) (**Figure 4A**). In the Δ*hofQ* and Δ*pilA::spe^r^*mutants, the expression of all six genes was either up- or downregulated compared to that in the wild-type strain, but the difference was not statistically significant for every gene (**Figure 4A**). The changes in the relative expression of five (Δ*hofQ*) and three (Δ*pilA::spe^r^*) genes were statistically significant (p = 0.0495, Mann‒Whitney U test). There was a high or extremely high correlation between the qPCR results and RNAseq results for five genes (Pearson’s R = 0.74–0.96, p < 0.05) (**Figure 4B**). The *nrfB* gene showed a lower yet moderate correlation (Pearson’s R = 0.59, p = 0.042) (**Figure 4B**). The observed correlations further validated the RNAseq results.

**Figure 4.**
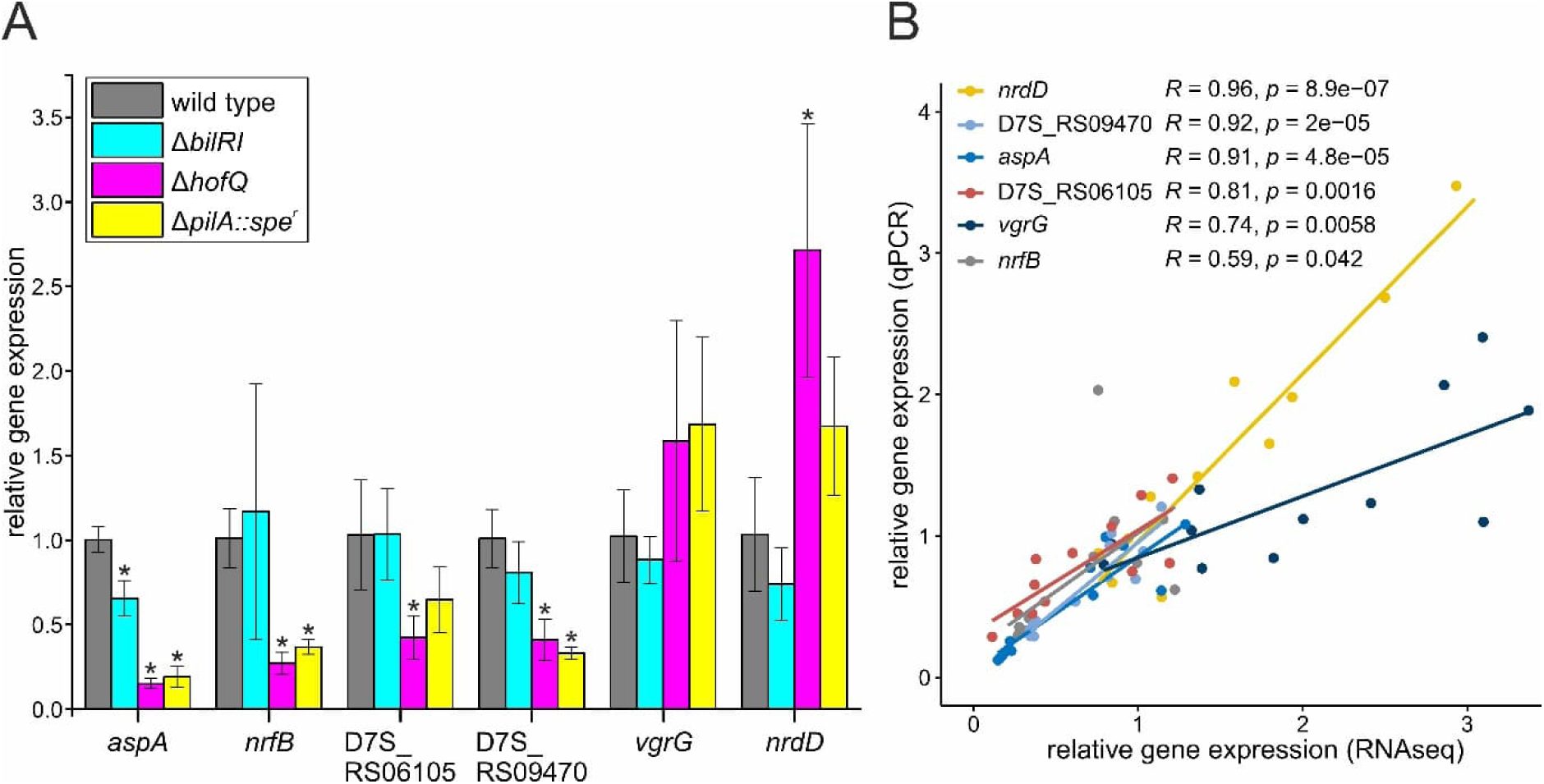
The RNA sequencing results were validated with qPCR. **A.** Relative expression levels of six selected genes were measured with qPCR. Five of the genes were significantly differentially expressed in the Δ*hofQ* mutant (p = 0.0495, Mann‒Whitney U test), three in the Δ*pilA::spe^r^* mutant (p = 0.0495, Mann‒Whitney U test) and one in the Δ*bilRI* mutant (p = 0.0495, Mann‒Whitney U test) compared to the wild-type strain. The qPCR results resembled the RNAseq results except for the significant downregulation of *aspA* in the Δ*bilRI* mutant, nonsignificant downregulation of D7S_RS06105 in the Δ*pilA::spe^r^* mutant and nonsignificant upregulation of *vgrG* in the Δ*hofQ* and Δ*pilA::spe^r^*mutants. **B.** There were moderate to extremely high correlations between the relative gene expression levels obtained by RNAseq and those obtained by qPCR (Pearson’s R = 0.59–0.96, p < 0.05).

There are eight highly similar copies of *vgrG* in the genome of *A. actinomycetemcomitans* D7S (NC_017846.2), although they vary in length (903–2223 nt). The qPCR primers used for amplification of *vgrG* at locus D7S_12360 also recognized other copies of the gene in the genome, which biased the results. This probably explains the differences in *vgrG* expression between the RNAseq and qPCR data. The RNAseq results showed a significant, greater than 2-fold increase in *vgrG* expression in Δ*hofQ* and Δ*pilA::spe^r^* compared to the wild-type strain, whereas the qPCR results indicated nonsignificant upregulation of less than 2-fold.

### The Δ*pilA::spe^r^* mutant contained a unique cluster of 15 genes

The Δ*pilA::spe^r^* mutant had more differentially regulated genes than the Δ*hofQ* mutant. However, among the upregulated genes in Δ*pilA::spe^r^* was a gene cluster that was not expressed in any of the other three strains, including the wild-type strain. This element comprises 15 genes (D7S_RS06500– D7S_RS06570), of which 10 were considered differentially expressed in the Δ*pilA::spe^r^* mutant. RNA sequencing revealed that none of the 15 genes were expressed in any other strain used in this study. PCR with primers targeting two of the genes was used to confirm the absence of this element in the other three strains (data not shown). The functions of most of the genes were unknown, but the known functions included host cell division inhibitor (D7S_RS06515), phage antirepressor (D7S_RS06525), transcriptional regulator (D7S_RS06550), phage regulatory protein (D7S_RS06555) and recombinase/integrase (D7S_RS06570). The protein products of genes D7S_RS06525 and D7S_RS06570 belong to COG category X (Mobilome: prophages, transposons). Moreover, the gene cluster is part of a genetic island in the D7S genome [48]. A nucleotide BLAST search using the genetic island region (1,310,166–1,331,249) as a template revealed that only two additional *A. actinomycetemcomitans* strains (624 and 14R) contain the same region (data not shown). A sequence nearly identical (23/24 nt) to the *Escherichia coli* prophage CP4-57 attachment site *attL* was also present in the region, overlapping the *ssrA* gene as observed in *E. coli* [49,50]. These observations suggest that the region could be a prophage [51].

### Genes associated with anaerobic respiration/metabolism were downregulated in the Δ*hofQ* mutant

Almost half of the downregulated genes (10 of 22) in the Δ*hofQ* mutant are associated with anaerobic respiration or anaerobic metabolism. Downregulated genes associated with anaerobic respiration were the periplasmic nitrite reductase operon *nrfABCD* (D7S_RS09090-09105) and those encoding the heme exporter protein CcmD (D7S_RS09050), c-type cytochrome biogenesis protein CcmI (D7S_RS09075), and cytochrome c membrane protein NapC (D7S_RS06105). Other genes associated with anaerobiosis were the aspartate ammonia-lyase gene *aspA* (D7S_RS05920), whose protein product catalyzes the conversion of aspartate to fumarate and is therefore necessary for growth under anaerobic conditions [52], and two genes whose protein products participate in anaerobic arginine catabolism (D7S_RS08420-08425).

Although several genes associated with anaerobic conditions were downregulated, two genes associated with anaerobic DNA synthesis were upregulated: the anaerobic ribonucleoside-triphosphate gene *nrdD* (D7S_RS03435) and its activator *nrdG* (D7S_RS03440) [53]. Moreover, *ybgE* (D7S_RS02890), which is located in the cytochrome bd operon upstream of the *cyd* genes, was downregulated in Δ*hofQ*. However, although cytochrome bd is typically activated at low oxygen levels in facultative anaerobes, the *ybgE* gene has been shown to be dispensable for oxidase activity [54].

In addition to the DNA synthesis genes *nrdD* and *nrdG*, three more genes associated with transcription or translation were regulated in the Δ*hofQ* mutant. While the transcriptional repressor *metJ* (D7S_RS04710) was downregulated, indicating activation of the methionine regulon [55], the IMP dehydrogenase gene *guaB* (D7S_RS10615) and ribosome maturation factor gene *rimP* (D7S_RS01570) were upregulated. IMP dehydrogenase participates in *de novo* synthesis of guanine nucleotides, and *rimP* is required for maturation of 30S ribosomal subunits. Moreover, a gene at locus D7S_RS03355 encoding a protein homologous to the *E. coli* transcription antiterminator CspE [56] was upregulated. This protein is also characterized as a cold shock protein, and interestingly, the heat shock-associated chaperones ClpB and DnaJ, along with the protease HslV, were downregulated.

Additionally, two putative virulence factors were upregulated in Δ*hofQ*: the type IV secretion system tip protein-encoding gene *vgrG* (D7S_RS12360) [57] and the γ-glutamyltransferase gene *ggt* (D7S_RS00965) [58]. Other downregulated genes were two CRISPR-associated genes (D7S_RS00920- 00925), the membrane lipid synthesis enzyme-encoding gene *fabA* (D7S_RS00590), and the gene encoding a membrane transport protein homologous to lipid transporters (D7S_RS09470). Four downregulated proteins had uncharacterized functions; however, one was a putative invasion gene upregulator (SirB2 family protein, D7S_RS00815). Additionally, the 6S RNA gene (*ssrS*, D7S_RS03800) was downregulated in the Δ*hofQ* mutant. Overall, the differentially regulated genes in the Δ*hofQ* mutant suggest that this mutant exhibits inhibition of anaerobic respiration and an increase in DNA and protein synthesis.

### Genes associated with anaerobic respiration/metabolism were downregulated in the Δ*pilA::spe^r^* mutant

Similar to the findings in the Δ*hofQ* mutant, several genes associated with anaerobic respiration were downregulated in the Δ*pilA::spe^r^* mutant. Altogether, eleven proteins associated with anaerobic respiration were downregulated: the proteins encoded by the periplasmic nitrite reductase operon *nrfABCD* (D7S_RS09090-09105), the heme exporter protein CcmD (D7S_RS09050), the proteins encoded by the periplasmic quinol-oxidizing system genes *napGH* (D7S_RS06090-06095), the cytochrome c subunit NapB (D7S_RS06100), the cytochrome c membrane protein NapC (D7S_RS06105), the cytochrome c peroxidase (D7S_RS11350) and the protein encoded by the dimethylsulfoxide reductase subunit b gene *dmsB* (D7S_RS06385). Some of these enzymes require molybdenum (Mo) as a cofactor [59], and accordingly, the molybdopterin synthase subunit gene *moaD* (D7S_RS00020) and molybdate-binding protein-encoding gene *modA* (D7S_RS01140) were also downregulated.

Interestingly, the expression of the malate dehydrogenase gene *mdh* (D7S_RS01080) was downregulated in the Δ*pilA::spe^r^* mutant. Malate dehydrogenase catalyzes the oxidation of malate to oxaloacetate under aerobic conditions but also catalyzes the reversible reaction under anaerobic conditions [60]. The iron-sulfur cluster insertion protein ErpA, which is essential for both aerobic and anaerobic respiration in *E. coli* [61], was also downregulated. Other downregulated enzymes associated with anaerobiosis were the aspartate ammonia-lyase *aspA* (D7S_RS05920); the L-asparaginase II *ansB* (D7S_RS08840), whose expression has been shown to increase under anaerobic conditions [62]; and the 3-keto-L-gulonate-6-phosphate decarboxylase UlaD (D7S_RS02685), which is a component of the anaerobic L-ascorbate fermentation pathway [63]. One additional downregulated protein was a YdcF family protein (D7S_RS05750) homologous to *E. coli* YdcF, which has been suggested to be involved in the anaerobic respiration pathway [64]. A BLAST search also revealed similarities to the conserved protein domain family member NfrG (COG4235), which is characterized as a cytochrome c-type biogenesis protein. However, the COG database search indicated that the YdcF family protein is a membrane biogenesis protein.

Although multiple genes associated with anaerobic respiration were downregulated, the expression of formate dehydrogenase-N subunits α (*fdnG*) and β (*fdxH*) was increased in the Δ*pilA::spe^r^* mutant. Formate dehydrogenase-N also participates in anaerobic respiration [65].

### Genes associated with stress were upregulated in the Δ*pilA::spe^r^* mutant

The expression of several stress response proteins was upregulated in the Δ*pilA::spe^r^* mutant. The RNA polymerase sigma factor RpoE, which is an extracytoplasmic stress factor [66], was upregulated along with its regulators, including the downstream gene *rseA* (D7S_RS04115), protease gene *rseP* (D7S_RS04270) and protease DegP homolog (D7S_RS01980) [67]. Additionally, the downstream genes *rseB* and *rseC* (D7S_RS04110–04105) were downregulated, although the change in its expression was just below the set threshold (2.0-fold change). Several RpoE regulon genes (studied in *E. coli* [68–70]) were upregulated in the Δ*pilA::spe^r^* mutant, such as those encoding the protease Lon (D7S_RS08780), chaperones DnaJ (D7S_RS09770) and DnaK (D7S_RS09775), and an uncharacterized YcbK family protein (D7S_RS10130). However, certain RpoE regulon genes were downregulated, including the L-asparaginase 2 gene *ansB* (D7S_RS08840), the gene encoding the CRISPR-associated helicase/endonuclease Cas3 (D7S_RS00925) and the N-acetylneuraminate anomerase gene *nanQ* (D7S_RS08015).

The heat shock factor RpoH, whose expression is positively regulated by RpoE [67], was also upregulated. The expression of several RpoH regulon proteins (studied in *E. coli* [71] and *P. aeruginosa* [72]) was also increased. These included the molecular chaperones GroEL (D7S_RS05905), GroES (D7S_RS05910), HtpG (D7S_RS06880), ClpB (D7S_RS06290), DnaJ (D7S_RS09770) and DnaK (D7S_RS09775); the RNA chaperone Hfq (D7S_RS10695); the proteases HslV (D7S_RS05320), HslU (D7S_RS05325) and Lon (D7S_RS08780); the tRNA dimethylallyltransferase MiaA (D7S_RS10700); and an SNF2-related transcriptional regulator homologous to *E. coli* RapA (HepA) (D7S_RS09960).

Two additional genes that were upregulated in the Δ*pilA::spe^r^* mutant are also associated with cellular stress. A gene at locus D7S_RS00715 encodes a Sel1-repeat family protein that can be activated upon cellular stress [73]. Another gene at locus D7S_RS10315 encodes a YoeB family toxin, which is a putative translation inhibitor and associated with thermal stress [74].

### Genes associated with cell membrane biogenesis were upregulated in the Δ*pilA::spe^r^* mutant

Ten genes associated with cell membrane synthesis and integrity or cell division were upregulated in the Δ*pilA::spe^r^* mutant, indicating a possible increase in the cell division rate in comparison to that of the wild-type strain. The ten genes included four involved in peptidoglycan synthesis and cell wall formation (*glmS* (D7S_RS04570)*, mltF* (D7S_RS08935), *murA* (D7S_RS09480), and a ligase (D7S_RS06850)), three Tol-Pal system genes (*tolA*, *tolB*, and *tolR* (D7S_RS02865–02875)), the gene encoding the lipid asymmetry maintenance protein MlaD (D7S_RS09500), the gene encoding an RlpA family protein (D7S_RS05505) and the gene encoding a ParA family protein (D7S_RS09725). Additionally, *lpxH* (D7S_RS09285), whose protein product is involved in the biosynthesis of lipid A, was upregulated.

### Differential regulation of virulence-associated genes

The only upregulated gene common to the Δ*hofQ* and Δ*pilA::spe^r^*mutants was *vgrG*, which encodes the type VI secretion system tip protein. More specifically, *vgrG* at locus D7S_RS12360 was upregulated. However, the qPCR results indicated moderate upregulation of *vgrG*, even though the primers targeted multiple *vgrG* copies in the *A. actinomycetemcomitans* D7S genome (see “qPCR validation of the RNAseq results”). The type VI secretion system is a putative virulence factor in pathogenic species such as *Acinetobacter baumannii* [57] and *Burkholderia* spp. [75].

Specific virulence-associated genes were downregulated in the Δ*pilA::spe^r^*mutant. Glycosyltransferase family 9 proteins (D7S_RS04980–04985) likely participate in LPS synthesis [76], whereas the toxin-activating lysine-acyltransferase, also called LtxC (D7S_RS02775), is required for activation of leukotoxin (encoded by *ltxA*) [77]. Further examination of the RNAseq results revealed that the *ltxA* gene (D7S_RS02770) was also downregulated, although the change in its expression (1.8- fold) was less than 2-fold, which was used as the threshold for a significant change in expression. The major fimbrial subunit Flp-1 (D7S_RS06730), required for nonspecific tight adherence in *A. actinomycetemcomitans* [78], was surprisingly downregulated, along with the quorum sensing gene *luxS* (D7S_RS06455). Both *flp-1* and *luxS* are required for formation of *A. actinomycetemcomitans* biofilms [78,79]. Additionally, the transformation regulatory gene *tfoX (sxy)* [22] was downregulated, indicating downregulation of competence-associated genes. However, except for the deleted *pilA* gene, the only downregulated competence gene was the gene encoding the type IV pilus transport protein PilC (D7S_RS07995), which belongs to the same cluster as *pilA* [21].

### Differential regulation of genes involved in carbohydrate or iron transport/metabolism in the Δ*pilA::spe^r^* mutant

In the Δ*pilA::spe^r^* mutant, several carbohydrate transport- or metabolism-associated genes were differentially regulated. Two phosphotransferase system (PTS) genes, which are located directly downstream and in the same direction as the gene encoding the 3-keto-L-gulonate-6-phosphate decarboxylase UlaD involved in the L-ascorbate fermentation pathway, were also downregulated. Based on homology, both subunit IIA (D7S_RS02680) and IIBC (D7S_RS02675) are ascorbate specific, which could explain their potential coregulatory relationship with UlaD. Other carbohydrate transport and metabolism genes were also downregulated, including genes involved in sugar transport (*mglB* (D7S_RS07120) and D7S_RS04410-04415), ribose metabolism (*rbsA* and *rbsD* (D7S_RS03335–03330)), and glycogen biosynthesis (*glgC* (D7S_RS00395)). Instead, galactitol-specific PTS transporter subunit IIC (D7S_RS12375) and two genes whose encoded proteins are involved in gluconate uptake and metabolism (D7S_RS10290–10295) were upregulated.

Along with carbohydrate transporters, other transporter proteins were also upregulated in the Δ*pilA::spe^r^* mutant. The increased expression of an iron chelate uptake ABC transporter protein (D7S_RS01000-01010) and TonB system transport protein (D7S_RS05280-05285) indicate enhanced iron uptake and can be linked with iron limitation [80]. However, the protein products of the genes at loci D7S_RS00260–00270 and D7S_RS04930 are involved in type I secretion, based on their homology to HlyD and RND family adaptors [81].

## Discussion

Interestingly, deletion of *hofQ* or *pilA* in the *A. actinomycetemcomitans* D7S strain led to downregulation of several genes associated with anaerobic respiration or anaerobic metabolism. Previous studies with *A. actinomycetemcomitans* have shown that genes associated with anaerobic respiration/metabolism are repressed by iron limitation and induced by *in vivo* growth (murine abscess model) and by catecholamine- and iron-induced activation of QseBC [80,82,83]. Because QseBC is required for efficient biofilm formation and virulence in *A. actinomycetemcomitans* [84], it is likely that the QseBC regulon contributes to virulence [83]. Induced anaerobic respiration/metabolism has thus been suggested to prime the cell population to inhabit anaerobic niches *in vivo* and increase virulence. Therefore, downregulation of genes associated with an anaerobic lifestyle in the Δ*hofQ* and Δ*pilA::spe^r^* mutants could indicate a decreased colonization ability and decreased virulence. Along with the downregulation of anaerobic respiration, iron uptake was upregulated in the Δ*pilA::spe^r^* mutant, indicating a link between respiration and iron acquisition similar to that found in the studies mentioned above.

The decreased colonization ability and virulence of Δ*pilA::spe^r^* was also implied by the downregulation of genes directly associated with biofilm formation and virulence. The major pilin subunit Flp-1, which belongs to the tight adherence (tad) locus, was downregulated in the Δ*pilA::spe^r^* mutant, along with autoinducer-2 (*luxS*), which mediates quorum sensing. The downregulation of these two genes indicates impaired biofilm formation because they are required for efficient formation of *A. actinomycetemcomitans* biofilms [78,79]. Neither Flp-1 nor *luxS* was differentially regulated in the Δ*hofQ* mutant; however, 6S RNA (*ssrS*) was downregulated, which could also indicate impaired biofilm formation [85]. Biofilm dispersion is mediated by dispersin B (*dspB* (D7S_RS07210)), which is induced by iron limitation [80]. The upregulation of genes involved in iron uptake observed in the Δ*pilA::spe^r^* mutant resembles the transcription profile during iron limitation. Thus, more in-depth analysis of the RNAseq data revealed that *dspB* was slightly yet significantly upregulated in the Δ*pilA::spe^r^*mutant (fold change +1.6). Although slight upregulation of *dspB* was also observed in the Δ*hofQ* mutant, the difference was not statistically significant. Overall, the observed downregulation of Flp-1 and *luxS* along with upregulation of *dspB* indicate that the Δ*pilA::spe^r^* mutant might not form as robust a biofilm as the wild-type strain. Additionally, *luxS* has been suggested to contribute to the virulence of *A. actinomycetemcomitans* by inducing leukotoxin expression [86]. Accordingly, the leukotoxin- activating gene *ltxC* was downregulated in the Δ*pilA::spe^r^* mutant, indicating decreased virulence. Moreover, PTS has been associated with biofilm formation in some pathogenic species [87–90]. Three PTS genes were differentially regulated in the Δ*pilA::spe^r^* mutant: two ascorbate-specific subunits were downregulated, whereas one galactitol-specific subunit was upregulated. However, it is unclear how the ascorbate- and galactitol-specific PTS enzymes affect biofilm formation, as previous studies have focused on the importance of glucose- and mannose-specific PTS enzymes.

Biofilm formation is essential for the virulence of many pathogenic species. Cells in biofilms exhibit physiological differences depending on which part of the biofilm structure they inhabit. This biofilm heterogeneity is the result of chemical gradients formed within the layers of the biofilm, which induce changes in gene expression leading to changes in protein production and metabolic activity [4,91]. Therefore, harvesting the whole biofilm population for transcriptional analysis results in an average of the physiological state of the cell population. Studies performed with *P. aeruginosa* show that cells residing in the top layers of a biofilm are typically more metabolically active and express genes required for growth and division [92]. Additionally, proteins encoded by stress response genes, including the alternative sigma factor RpoH and its regulon, are highly abundant in the top layers of the biofilm compared to the bottom layers [92]. The stress response was induced in the Δ*pilA::spe^r^* mutant, which could indicate that a greater proportion of the biofilm population is considered the “top layer”, meaning that the biofilm formed by the Δ*pilA::spe^r^* mutant is not as thick or robust as that formed by the wild-type strain. Although only seven protein-coding genes were upregulated in the Δ*hofQ* mutant, all seven were associated with transcription/translation, thus indicating increased metabolic activity compared to the wild-type strain. Therefore, the observed changes in gene expression could indicate changes in biofilm formation in the Δ*hofQ* and Δ*pilA::spe^r^* mutants.

Along with RpoH, the alternative sigma factor RpoE was also upregulated in the Δ*pilA::spe^r^* mutant, implying induction of the stress response. The RpoE regulon (studied in *E. coli* [68–70]) includes proteins involved in virulence, such as those participating in assembly and maintenance of LPS. However, increased expression of such genes, except for one gene, *lpxH*, involved in lipid A biosynthesis, was not observed. In contrast, putative LPS synthesis genes, along with certain virulence genes, were downregulated, indicating decreased virulence. However, the outer membrane of gram-negative bacteria is also regulated by RpoE in response to environmental changes, and several genes associated with membrane biogenesis were upregulated in the Δ*pilA::spe^r^* mutant.

Overall, the changes in gene expression resulting from deletion of either *hofQ* or *pilA* indicate decreased biofilm formation. These results are consistent with previous observations of decreased biofilm formation of *A. actinomycetemcomitans* D7S Δ*hofQ* [17]. Moreover, deletion of *bilRI* did not result in changes in gene expression, consistent with previous results showing that the biofilm formation of Δ*bilRI* does not differ from that of the wild-type strain [16]. The biofilm formation of Δ*pilA::spe^r^*has not been studied in detail, but the deletion mutant exhibits a colony morphology similar to that of the wild-type strain [21]. Although *bilRI* has been shown to decrease the transformation efficiency [23], deletion of the gene does not result in a completely nontransformable strain, as observed with deletion of *hofQ* in this study and with deletion of *pilA* previously [21]. Furthermore, unlike the other two proteins, BilRI does not bind to USS-dsDNA [23]. Therefore, it is possible that the expression of genes essential for competence, such as *hofQ* and *pilA*, is required for the virulence of the naturally competent *A. actinomycetemcomitans* strain. However, further studies are needed to confirm this hypothesis.

Although multiple genes were differentially regulated in the Δ*pilA::spe^r^* deletion mutant, it is not clear whether all changes occurred solely as a result of the deletion. The presence of a prophage-like element in the genome of the Δ*pilA::spe^r^* mutant raises the question of which changes in gene expression resulted from deletion of *pilA* rather than translation of the prophage-like genes. Although all the deletion mutants were generated from *A. actinomycetemcomitans* strain D7S, only Δ*bilRI* and Δ*hofQ* were generated using the same wild-type isolate that was used in this study. In contrast, the Δ*pilA::spe^r^* mutant was not generated in our laboratory but was received as a kind gift from Prof. Chen [21]. It is possible that the prophage was excised spontaneously, which could explain its absence in the other strains in this study. The biofilm formation and virulence of many species can be affected by the presence of an active prophage element in their genome [93–95]. Future studies must be performed to investigate the effect of prophage-like elements on biofilm formation and global gene expression in *A. actinomycetemcomitans*. Moreover, the effect of deletion of *pilA* on the gene expression profile of *A. actinomycetemcomitans* D7S must be confirmed using wild-type and mutant strains that either do or do not possess the prophage-like element. However, the Δ*pilA::spe^r^* mutant shared common differentially regulated genes with the Δ*hofQ* mutant, which does not possess the prophage-like element—most notably, genes associated with anaerobic respiration. This observation suggests that deletion of the competence gene *pilA* indeed affects global gene expression in naturally competent *A. actinomycetemcomitans* D7S.

## Acknowledgments

We thank Finnish Functional Genomics Centre supported by University of Turku, Åbo Akademi University and Biocenter Finland. This work was financially supported by the Academy of Finland under grant number 322817 and the University of Turku Graduate School.

## Notes

### Competing Interest Statement

The authors have declared no competing interest.

