## Supplemental Tables 1-4 for "Deletion of competence genes represses expression of genes associated with anaerobic respiration/metabolism in *Aggregatibacter actinomycetemcomitans*"

Tykistökatu 6A

20014 University of Turku, FINLAND

### Supplemental tables S1-S4

The supplemental tables present the differentially regulated genes in *A. actinomycetemcomitans* Δ*hofQ* and Δ*pilA::spe*$^{r}$. The tables include genes whose expression had changed at least 2-fold compared to the wild type strain and had a Benjamini–Hochberg adjusted p value of < 0.05.

Table S1: Upregulated genes in *A. actinomycetemcomitans* Δ*hofQ*.

| locus tag | gene | product | COG cat. | fold change | p-value |
|---|---|---|---|---|---|
| D7S_RS03440 | *nrdG* | anaerobic ribonucleoside-triphosphate reductase-activating protein | H | 3.20 | 4.281e-10 |
| D7S_RS03435 | *nrdD* | anaerobic ribonucleoside-triphosphate reductase | F | 2.51 | 1.940e-05 |
| D7S_RS12360 | *vgrG* | type VI secretion system tip protein VgrG | NA | 2.51 | 4.858e-03 |
| D7S_RS00965 | *ggt* | gamma-glutamyltransferase | E | 2.27 | 4.196e-04 |
| D7S_RS10615 | *guaB* | IMP dehydrogenase | F, K | 2.14 | 1.078e-03 |
| D7S_RS03355 | *NA* | cold-shock protein | K | 2.10 | 3.748e-06 |
| D7S_RS00135 | *NA* | NA: pseudoprotein | NA | 2.03 | 3.385e-02 |
| D7S_RS01570 | *rimP* | ribosome maturation factor RimP | J | 2.00 | 5.318e-05 |

Table S2: Downregulated genes in *A. actinomycetemcomitans* Δ*hofQ*.

| locus tag | gene | product | COG cat. | fold change | p-value |
|---|---|---|---|---|---|
| D7S_RS05070 | *pilQ* | type IV pilus secretin PilQ (= HofQ) | U | -138.14 | 2.872e-27 |
| D7S_RS05920 | *aspA* | aspartate ammonia-lyase | E | -5.82 | 3.088e-30 |
| D7S_RS09095 | *nrfB* | cytochrome c nitrite reductase pentaheme subunit | NA | -3.86 | 7.917e-22 |
| D7S_RS09090 | *nrfA* | ammonia-forming nitrite reductase cytochrome c552 subunit | NA | -3.78 | 1.929e-24 |
| D7S_RS06105 | *NA* | NapC/NirT family cytochrome c | C | -3.66 | 3.126e-04 |
| D7S_RS09100 | *nrfC* | cytochrome c nitrite reductase Fe-S protein | C | -3.41 | 5.995e-11 |
| D7S_RS09105 | *nrfD* | cytochrome c nitrite reductase subunit NrfD | P | -3.29 | 4.818e-11 |
| D7S_RS08360 | *NA* | NA: pseudoprotein | NA | -2.64 | 1.402e-05 |
| D7S_RS09050 | *ccmD* | heme exporter protein CcmD | U | -2.60 | 2.624e-05 |
| D7S_RS00815 | *NA* | SirB2 family protein | S | -2.48 | 1.245e-02 |
| D7S_RS11365 | *NA* | DUF302 domain-containing protein | S | -2.39 | 1.093e-07 |
| D7S_RS05895 | *NA* | tRNA-Pro | NA | -2.35 | 3.424e-03 |
| D7S_RS06290 | *clpB* | ATP-dependent chaperone ClpB | O | -2.31 | 8.543e-08 |
| D7S_RS08420 | *NA* | ornithine carbamoyltransferase | E | -2.23 | 2.770e-08 |
| D7S_RS09470 | *NA* | outer membrane protein transport protein | I | -2.19 | 8.920e-05 |
| D7S_RS00920 | *cas5c* | type I-C CRISPR-associated protein Cas5c | NA | -2.17 | 1.272e-08 |
| D7S_RS05320 | *hslV* | ATP-dependent protease subunit HslV | O | -2.17 | 2.717e-04 |
| D7S_RS03800 | *ssrS* | 6S RNA | NA | -2.13 | 7.568e-05 |
| D7S_RS01685 | *NA* | YacL family protein | S | -2.13 | 1.000e-02 |
| D7S_RS09075 | *ccmI* | c-type cytochrome biogenesis protein CcmI | O, C | -2.08 | 4.748e-08 |
| D7S_RS00590 | *fabA* | bifunctional 3-hydroxydecanoyl-ACP dehydratase/trans-2-decenoyl-ACP isomerase | I | -2.06 | 1.402e-08 |
| D7S_RS00925 | *NA* | CRISPR-associated helicase/endonuclease Cas3 | V | -2.06 | 7.666e-03 |
| D7S_RS02890 | *ybgE* | cyd operon protein YbgE | S | -2.04 | 2.421e-05 |
| D7S_RS08425 | *arcC* | carbamate kinase | E | -2.03 | 9.615e-08 |
| D7S_RS09770 | *dnaJ* | molecular chaperone DnaJ | K, E | -2.01 | 1.402e-05 |
| D7S_RS04710 | *metJ* | met regulon transcriptional regulator MetJ | O | -2.01 | 4.531e-03 |

Table S3: Upregulated genes in *A. actinomycetemcomitans* Δ*pilA::spe*$^{r}$.

| locus tag | gene | product | COG cat. | fold change | p-value |
|---|---|---|---|---|---|

| | | | | | |
|---|---|---|---|---|---|
| D7S_RS06515 | *NA* | host cell division inhibitor Icd-like protein | NA | 27175.14 | 1.735e-31 |
| D7S_RS06570 | *NA* | tyrosine-type recombinase/integrase | L, X | 5148.73 | 2.998e-22 |
| D7S_RS06510 | *NA* | hypothetical protein | NA | 1520.15 | 2.231e-16 |
| D7S_RS06500 | *NA* | hypothetical protein | NA | 1398.83 | 7.099e-16 |
| D7S_RS06560 | *NA* | MULTISPECIES: hypothetical protein | NA | 1120.56 | 3.557e-15 |
| D7S_RS06565 | *NA* | MULTISPECIES: hypothetical protein | NA | 849.22 | 5.190e-14 |
| D7S_RS06520 | *NA* | MULTISPECIES: winged helix-turn-helix domain-containing protein | NA | 265.03 | 2.657e-09 |
| D7S_RS06505 | *NA* | hypothetical protein | NA | 225.97 | 8.621e-09 |
| D7S_RS06525 | *NA* | MULTISPECIES: phage antirepressor Ant | X | 58.08 | 1.890e-04 |
| D7S_RS06545 | *NA* | hypothetical protein | NA | 58.08 | 1.939e-04 |
| D7S_RS12405 | *NA* | DUF1439 domain-containing protein | NA | 5.028 | 4.697e-17 |
| D7S_RS10700 | *miaA* | tRNA (adenosine(37)-N6)-dimethylallyltransferase MiaA | J | 4.79 | 1.141e-24 |
| D7S_RS12410 | *NA* | DUF1439 domain-containing protein | NA | 4.29 | 1.300e-11 |
| D7S_RS09775 | *dnaK* | molecular chaperone DnaK | O | 4.26 | 1.815e-10 |
| D7S_RS10625 | *NA* | glycine zipper 2TM domain-containing protein | NA | 4.08 | 6.598e-13 |
| D7S_RS05320 | *hslV* | ATP-dependent protease subunit HslV | O | 3.71 | 1.540e-09 |
| D7S_RS00715 | *NA* | sel1 repeat family protein | NA | 3.58 | 3.223e-19 |
| D7S_RS00270 | *NA* | HlyD family efflux transporter periplasmic adaptor subunit | V | 3.43 | 1.308e-14 |
| D7S_RS00720 | *NA* | YwiC-like family protein | NA | 3.34 | 5.474e-15 |
| D7S_RS12355 | *NA* | NA | NA | 3.34 | 5.251e-03 |
| D7S_RS06290 | *clpB* | ATP-dependent chaperone ClpB | O | 3.32 | 8.022e-07 |
| D7S_RS12255 | *NA* | hypothetical protein | NA | 3.18 | 1.099e-09 |
| D7S_RS10695 | *hfq* | RNA chaperone Hfq | T | 3.14 | 3.385e-20 |
| D7S_RS12360 | *vgrG* | type VI secretion system tip protein VgrG | NA | 3.07 | 3.315e-05 |
| D7S_RS07240 | *surE* | 5\\'/3\\'-nucleotidase SurE | L | 3.05 | 3.327e-14 |
| D7S_RS11085 | *NA* | hypothetical protein | I | 3.03 | 2.833e-16 |
| D7S_RS03760 | *NA* | NAD(P)H-dependent oxidoreductase | C | 2.97 | 2.403e-15 |
| D7S_RS06295 | *NA* | helix-hairpin-helix domain-containing protein | NA | 2.93 | 9.162e-09 |
| D7S_RS09770 | *dnaJ* | molecular chaperone DnaJ | O | 2.89 | 8.073e-11 |
| D7S_RS00265 | *rbbA* | ribosome-associated ATPase/putative transporter RbbA | V | 2.89 | 6.725e-09 |
| D7S_RS09300 | *NA* | aromatic amino acid transporter | E | 2.87 | 1.540e-07 |
| D7S_RS04570 | *glmS* | glutamine--fructose-6-phosphate transaminase (isomerizing) | M | 2.73 | 4.436e-09 |
| D7S_RS00005 | *NA* | NA | NA | 2.73 | 7.104e-06 |
| D7S_RS02110 | *NA* | integrase core domain-containing protein | NA | 2.71 | 2.270e-02 |
| D7S_RS02120 | *NA* | IS3 family transposase | NA | 2.68 | 1.677e-07 |
| D7S_RS04120 | *rpoE* | RNA polymerase sigma factor RpoE | K | 2.66 | 1.529e-03 |
| D7S_RS00135 | *NA* | NA | NA | 2.62 | 7.683e-08 |
| D7S_RS06850 | *NA* | D-alanine--D-alanine ligase | M, R | 2.60 | 2.777e-11 |
| D7S_RS02490 | *queD* | 6-carboxytetrahydropterin synthase QueD | H | 2.58 | 3.731e-09 |
| D7S_RS06435 | *NA* | PRD domain-containing protein | NA | 2.57 | 4.434e-13 |
| D7S_RS09450 | *glnA* | type I glutamate--ammonia ligase | E | 2.57 | 5.779e-07 |
| D7S_RS10260 | *fdxH* | formate dehydrogenase subunit beta | NA | 2.51 | 9.281e-06 |
| D7S_RS06880 | *htpG* | molecular chaperone HtpG | O | 2.51 | 1.867e-04 |
| D7S_RS01005 | *NA* | iron chelate uptake ABC transporter family permease subunit | P | 2.51 | 4.612e-03 |
| D7S_RS00260 | *NA* | ABC transporter permease | V | 2.43 | 5.412e-09 |
| D7S_RS09315 | *epmB* | EF-P beta-lysylation protein EpmB | NA | 2.40 | 6.849e-10 |
| D7S_RS04565 | *NA* | DeoR/GlpR family DNA-binding transcription regulator | K, G | 2.38 | 1.155e-11 |
| D7S_RS06060 | *rpoH* | RNA polymerase sigma factor RpoH | K | 2.38 | 1.836e-07 |
| D7S_RS02875 | *tolR* | colicin uptake protein TolR | U | 2.36 | 1.046e-06 |
| D7S_RS08780 | *lon* | endopeptidase La | NA | 2.36 | 1.889e-06 |
| D7S_RS01960 | *NA* | YegP family protein | S | 2.36 | 1.025e-04 |
| D7S_RS05905 | *groL* | chaperonin GroEL | O | 2.35 | 1.188e-04 |
| D7S_RS02870 | *tolA* | cell envelope integrity protein TolA | M | 2.33 | 2.063e-08 |
| D7S_RS04115 | *NA* | sigma-E factor negative regulatory protein | T | 2.33 | 1.498e-06 |
| D7S_RS01980 | *NA* | Do family serine endopeptidase | O | 2.31 | 9.719e-09 |

| | | | | | |
|---|---|---|---|---|---|
| D7S_RS07245 | *truD* | tRNA pseudouridine(13) synthase TruD | J | 2.31 | 2.199e-06 |
| D7S_RS09960 | *NA* | SNF2-related protein | K, L | 2.28 | 2.612e-04 |
| D7S_RS10130 | *NA* | YcbK family protein | S | 2.27 | 1.630e-08 |
| D7S_RS05040 | *NA* | 23S rRNA (adenine(2030)-N(6))-methyltransferase RlmJ | J | 2.23 | 1.453e-04 |
| D7S_RS09725 | *NA* | AAA family ATPase | NA | 2.22 | 2.314e-07 |
| D7S_RS02865 | *tolB* | Tol-Pal system beta propeller repeat protein TolB | U | 2.22 | 3.978e-07 |
| D7S_RS05325 | *hslU* | HslU--HslV peptidase ATPase subunit | O | 2.22 | 2.826e-06 |
| D7S_RS04930 | *NA* | efflux RND transporter periplasmic adaptor subunit | V, M | 2.22 | 1.521e-02 |
| D7S_RS02290 | *NA* | NA | NA | 2.22 | 1.842e-02 |
| D7S_RS10295 | *NA* | GntP family permease | G | 2.20 | 3.705e-05 |
| D7S_RS02495 | *NA* | 7-carboxy-7-deazaguanine synthase QueE | H | 2.20 | 1.872e-04 |
| D7S_RS01000 | *NA* | iron chelate uptake ABC transporter family permease subunit | P | 2.19 | 6.674e-05 |
| D7S_RS04540 | *NA* | hypothetical protein | NA | 2.19 | 7.090e-03 |
| D7S_RS09285 | *lpxH* | UDP-2,3-diacylglucosamine diphosphatase | M | 2.16 | 1.815e-10 |
| D7S_RS01010 | *NA* | ATP-binding cassette domain-containing protein | P | 2.16 | 1.828e-04 |
| D7S_RS10970 | *NA* | NA | NA | 2.16 | 1.162e-02 |
| D7S_RS12195 | *NA* | hypothetical protein | NA | 2.11 | 3.015e-03 |
| D7S_RS09500 | *mlaD* | outer membrane lipid asymmetry maintenance protein MlaD | M | 2.10 | 1.260e-04 |
| D7S_RS05280 | *exbD* | TonB system transport protein ExbD | U | 2.09 | 7.683e-08 |
| D7S_RS09995 | *cas8c* | NA | NA | 2.09 | 1.016e-05 |
| D7S_RS05505 | *NA* | septal ring lytic transglycosylase RlpA family protein | M | 2.07 | 1.157e-07 |
| D7S_RS11015 | *NA* | alcohol dehydrogenase catalytic domain-containing protein | E, R | 2.07 | 2.820e-06 |
| D7S_RS07200 | *pepB* | aminopeptidase PepB | E | 2.07 | 1.364e-05 |
| D7S_RS05285 | *NA* | energy transducer TonB | M | 2.07 | 1.130e-04 |
| D7S_RS08935 | *mltF* | membrane-bound lytic murein transglycosylase MltF | T, M | 2.06 | 1.105e-05 |
| D7S_RS10265 | *fdnG* | formate dehydrogenase-N subunit alpha | NA | 2.04 | 1.128e-06 |
| D7S_RS09480 | *murA* | UDP-N-acetylglucosamine 1-carboxyvinyltransferase | M | 2.04 | 1.260e-04 |
| D7S_RS12375 | *NA* | PTS galactitol transporter subunit IIC | NA | 2.04 | 6.502e-04 |
| D7S_RS04545 | *nudC* | NA | NA | 2.04 | 1.086e-03 |
| D7S_RS10315 | *NA* | Txe/YoeB family addiction module toxin | V | 2.03 | 3.749e-06 |
| D7S_RS04270 | *rseP* | sigma E protease regulator RseP | K, O | 2.01 | 5.524e-06 |
| D7S_RS05910 | *NA* | co-chaperone GroES | O | 2.00 | 9.236e-04 |
| D7S_RS10290 | *NA* | gluconokinase | G | 2.00 | 1.128e-03 |

Table S4: Upregulated genes in *A. actinomycetemcomitans ΔpilA::spe*$^r$.

| locus tag | gene | product | COG cat. | fold change | p-value |
|---|---|---|---|---|---|
| D7S_RS08005 | *NA* | prepilin-type N-terminal cleavage/methylation domain-containing protein (= PilA) | N, W | 404.50 | 2.275e-17 |
| D7S_RS06495 | *NA* | integrase | NA | 9.65 | 1.559e-11 |
| D7S_RS05920 | *aspA* | aspartate ammonia-lyase | E | 4.99 | 1.753e-21 |
| D7S_RS08375 | *NA* | ribosome alternative rescue factor ArfA | J | 4.59 | 2.474e-14 |
| D7S_RS11455 | *NA* | hypothetical protein | P | 4.53 | 2.836e-03 |
| D7S_RS09090 | *nrfA* | ammonia-forming nitrite reductase cytochrome c552 subunit | NA | 3.53 | 4.989e-24 |
| D7S_RS09095 | *nrfB* | cytochrome c nitrite reductase pentaheme subunit | NA | 3.27 | 1.873e-17 |
| D7S_RS01140 | *modA* | molybdate ABC transporter substrate-binding protein | P | 3.10 | 3.216e-15 |

| | | | | | |
|---|---|---|---|---|---|
| D7S_RS04410 | *NA* | sugar ABC transporter substrate-binding protein | G | 3.10 | 3.583e-05 |
| D7S_RS09100 | *nrfC* | cytochrome c nitrite reductase Fe-S protein | C | 3.01 | 3.685e-11 |
| D7S_RS02675 | *NA* | PTS ascorbate-specific subunit IIBC | G | 2.97 | 2.682e-14 |
| D7S_RS00920 | *cas5c* | type I-C CRISPR-associated protein Cas5c | NA | 2.89 | 1.208e-10 |
| D7S_RS02680 | *NA* | PTS sugar transporter subunit IIA | T, G | 2.87 | 6.044e-16 |
| D7S_RS09470 | *NA* | outer membrane protein transport protein | I | 2.87 | 5.715e-06 |
| D7S_RS02340 | *NA* | YeeE/YedE thiosulfate transporter family protein | NA | 2.85 | 2.152e-06 |
| D7S_RS05895 | *NA* | NA | NA | 2.77 | 1.146e-04 |
| D7S_RS09105 | *nrfD* | cytochrome c nitrite reductase subunit NrfD | P | 2.75 | 3.665e-11 |
| D7S_RS06105 | *NA* | NapC/NirT family cytochrome c | C | 2.73 | 2.870e-07 |
| D7S_RS00510 | *pepE* | dipeptidase PepE | E | 2.71 | 8.126e-10 |
| D7S_RS07070 | *NA* | NA | NA | 2.68 | 2.548e-04 |
| D7S_RS08360 | *NA* | NA | NA | 2.68 | 4.899e-06 |
| D7S_RS00605 | *cspD* | cold shock domain-containing protein CspD | K | 2.60 | 1.202e-05 |
| D7S_RS07120 | *mglB* | galactose/glucose ABC transporter substrate-binding protein MglB | G | 2.55 | 8.378e-08 |
| D7S_RS08840 | *ansB* | L-asparaginase 2 | J, E | 2.53 | 3.579e-13 |
| D7S_RS05750 | *NA* | YdcF family protein | M | 2.51 | 2.221e-15 |
| D7S_RS06100 | *NA* | nitrate reductase cytochrome c-type subunit | C, P | 2.51 | 3.206e-05 |
| D7S_RS06790 | *NA* | carbon starvation protein A | C, E | 2.50 | 4.407e-14 |
| D7S_RS09050 | *ccmD* | heme exporter protein CcmD | U | 2.45 | 5.083e-06 |
| D7S_RS00925 | *NA* | CRISPR-associated helicase/endonuclease Cas3 | V | 2.43 | 1.390e-10 |
| D7S_RS04985 | *NA* | glycosyltransferase family 9 protein | M | 2.43 | 1.143e-07 |
| D7S_RS12025 | *NA* | hypothetical protein | NA | 2.41 | 1.685e-07 |
| D7S_RS11350 | *NA* | cytochrome-c peroxidase | O | 2.40 | 1.536e-10 |
| D7S_RS03330 | *rbsD* | D-ribose pyranase | G | 2.38 | 3.583e-05 |
| D7S_RS12020 | *NA* | hypothetical protein | NA | 2.35 | 3.041e-10 |
| D7S_RS04825 | *NA* | YggL family protein | S | 2.33 | 3.005e-05 |
| D7S_RS06055 | *NA* | TfoX/Sxy family DNA transformation protein | K | 2.30 | 2.932e-10 |
| D7S_RS08430 | *NA* | YfcC family protein | R | 2.30 | 2.001e-06 |
| D7S_RS05760 | *deoC* | deoxyribose-phosphate aldolase | F | 2.28 | 2.105e-09 |
| D7S_RS10950 | *NA* | fructosamine kinase family protein | G | 2.28 | 1.001e-06 |
| D7S_RS02650 | *NA* | hypothetical protein | NA | 2.28 | 1.188e-04 |
| D7S_RS04980 | *NA* | glycosyltransferase family 9 protein | M | 2.22 | 6.287e-06 |
| D7S_RS07305 | *NA* | NA | NA | 2.22 | 1.077e-02 |
| D7S_RS06095 | *napH* | quinol dehydrogenase ferredoxin subunit NapH | C | 2.20 | 6.313e-04 |
| D7S_RS01080 | *mdh* | malate dehydrogenase | C | 2.19 | 3.846e-09 |
| D7S_RS05690 | *atpE* | MULTISPECIES: F0F1 ATP synthase subunit C | C | 2.19 | 1.309e-06 |
| D7S_RS02775 | *NA* | toxin-activating lysine-acyltransferase | O | 2.17 | 1.702e-03 |
| D7S_RS02355 | *NA* | hypothetical protein | NA | 2.17 | 2.471e-02 |
| D7S_RS00915 | *cas8c* | type I-C CRISPR-associated protein Cas8c/Csd1 | V | 2.16 | 5.360e-07 |
| D7S_RS02230 | *NA* | TIGR04211 family SH3 domain-containing protein | R | 2.14 | 1.053e-06 |
| D7S_RS08380 | *NA* | DUF5377 family protein | NA | 2.14 | 2.339e-05 |
| D7S_RS00020 | *moaD* | molybdopterin synthase sulfur carrier subunit | H | 2.13 | 1.247e-03 |
| D7S_RS06425 | *NA* | NA | NA | 2.13 | 1.183e-02 |
| D7S_RS06455 | *luxS* | S-ribosylhomocysteine lyase | T | 2.10 | 3.481e-10 |
| D7S_RS03335 | *rbsA* | ribose ABC transporter ATP-binding protein RbsA | G | 2.10 | 3.050e-07 |
| D7S_RS04415 | *NA* | sugar ABC transporter ATP-binding protein | G | 2.10 | 1.173e-05 |
| D7S_RS12125 | *NA* | DUF262 domain-containing protein | NA | 2.10 | 2.605e-04 |
| D7S_RS07995 | *NA* | type II secretion system F family protein | N, W, U | 2.08 | 2.850e-02 |
| D7S_RS01680 | *erpA* | iron-sulfur cluster insertion protein ErpA | O | 2.07 | 4.643e-05 |
| D7S_RS01200 | *NA* | hypothetical protein | NA | 2.07 | 7.229e-05 |
| D7S_RS07150 | *NA* | disulfide bond formation protein B | O | 2.06 | 1.182e-06 |
| D7S_RS06090 | *napG* | ferredoxin-type protein NapG | C | 2.04 | 5.427e-03 |

| | | | | | |
|---|---|---|---|---|---|
| D7S_RS02685 | *NA* | 3-keto-L-gulonate-6-phosphate decarboxylase UlaD | G | 2.01 | 1.213e-07 |
| D7S_RS06385 | *dmsB* | dimethylsulfoxide reductase subunit B | C | 2.01 | 3.315e-05 |
| D7S_RS06730 | *NA* | Flp family type IVb pilin | W, U | 2.01 | 1.086e-03 |
| D7S_RS00395 | *glgC* | glucose-1-phosphate adenylyltransferase | G | 2.01 | 7.198e-03 |
| D7S_RS08015 | *nanQ* | N-acetylneuraminate anomerase | G | 2.00 | 3.050e-07 |
